# Analysis of rhizosphere bacterial diversity of *Angelica dahurica* var. *formosana* based on different experimental sites and varieties (strains)

**DOI:** 10.1101/2021.06.24.449857

**Authors:** Meiyan Jiang, Fei Yao, Yunshu Yang, Yang Zhou, Kai Hou, Yinyin Chen, Dongju Feng, Wei Wu

**Affiliations:** Agronomy College, Sichuan Agricultural University, Wenjiang 611130, Chengdu Sichuan, P. R. China

**Keywords:** *Angelica dahurica*, rhizosphere, bacteria, yield, quality

## Abstract

Only with good producing areas and good germplasm can good medicinal materials be produced. In this study, in order to explore the effects of soil environment and germplasm on yield and quality of *Angelica dahurica* var. *formosana* from microorganism level, Illumina MiSeq was used to study the rhizosphere bacterial diversity of *A. dahurica* var. *formosana* based on different origin and varieties (strains). The results showed that the bacterial community of *A. dahurica* var. *formosana* was stable and conserved to a certain extent, and the bacteria of *Proteobacteria* and *Firmicutes* might play an important role in improving the yield and quality of *A. dahurica* var. *formosana*. Soil variables were the key factors affecting the rhizosphere bacterial community composition of *A. dahurica* var. *formosana*. These results are of great significance for further understanding the growth promotion mechanism of beneficial rhizosphere bacteria, reducing fertilizer application, expanding planting area, improving soil environment, and improving yield and quality.

**Importance:** Rhizosphere microorganisms play an important role in plant growth and accumulation of secondary metabolites. In this study, it was found for the first time that the rhizosphere bacterial community of *A. dahurica* var. *formosana* was relatively stable and conserved which could adapt to different environmental changes and resist the interference of external factors to a certain extent, providing a theoretical basis for distant planting and promotion of planting. The bacterial taxa of *Proteobacteria* and *Firmicutes* might play an important role in improving the yield and quality of *A. dahurica* var. *formosana*. This provided a theoretical basis for the exploitation and utilization of rhizosphere microbial resources of *A. dahurica* var. *formosana* to artificially control the soil environment and provided a new idea for increasing yield and quality of *A. dahurica* var. *formosana*.

## Introduce

Rhizosphere is a specific microecosystem of soil-root-microorganism interaction, which is formed by different plant, soil and environmental conditions. Meanwhile, the rhizosphere microbial community is also considered to be the second genome of plants (3). Plant roots can customize their rhizosphere microbial community by producing secretions, which in turn can affect plant growth through mineral nutrient absorption, antagonism, nutrition and spatial site competition, soil improvement and plant induced system resistance generation (8). Meanwhile, the rhizosphere microbial community construction is affected by a variety of biotic and abiotic factors. The specific factor that plays a decisive role varies by the soil type and plant species. The stronger the environmental screening effect of which factor is, the greater the relative contribution to the microbial community construction may be (25). Therefore, understanding the key factors affecting the composition and construction of beneficial bacterial communities in the rhizosphere may be a sustainable and effective way to actively manage soil microbial communities, improve soil conditions, reduce the application of pesticides and fertilizers, and promote plant growth (13).

*Angelica dahurica* var. *formosana* (known as Baizhi in Chinese) produced in Sichuan province is a famous Genuine Medicines, particularly in the Suining region of Sichuan province. It does not only serve for medicinal use in curing colds, headaches, nasal congestion, runny nose, toothache, but also possesses high commercial value for its wide adoption in the production of food, health care products, spices, skin care, beauty and other aspects. Modern pharmacological studies have found that coumarins, as the main index components of *A. dahurica* var. *formosana*, have antiinfammatory (12), anti-bacterial (33), vasodilator (39), anti-cancer (19), antiviral and antioxidant effects (2), and are regarded as one of the most important indicators to measure the quality of *A. dahurica* var. *formosana*. The *A. dahurica* var. *formosana* produced in Suining of Sichuan Province has a long history of cultivation with high yield and quality. However, some conditions during planting *A. dahurica* var. *formosana*, like early bolting, shrinking production areas, and abuse of pesticides and fertilizers are leading to its decline of the total yield of in recent years.

In order to solve the problem of production decline while ensuring quality, we could look for methods from a completely new angle, the beneficial bacterial community in the rhizosphere. Explore the secret of high yield and quality from rhizosphere from good origin to excavate the beneficial bacterial community structure. Meanwhile, plants with better germplasm will recruit more suitable microbe to promote their own growth. The development and utilization of rhizosphere bacteria resources of *A. dahurica* var. *formosana* will be a good method to improve the soil environment, reduce the application of fertilizer to a certain extent, and expand the planting region.

Therefore, this study aims to characterize bacterial communities in the rhizospheric soils of *A. dahurica* var. *formosana* from different experimental sites or varieties (strains) under the same culture management method by using molecular methods. This study may be a cornerstone of further research on biological factors to improve the quality of *A. dahurica* var. *formosana*. The objects of the present study are (i) to characterize the rhizospheric bacterial community in the samples of *A. dahurica* var. *formosana* from different sites or varieties (strains); (ii) to predict the related bacterial community closely related to yield and quality of *A. dahurica* var. *formosana*; (iii) to determine the contribution of experimental sites and its own varieties (strains) differences to the construction of beneficial bacterial community, and confirm the importance of good origins and good germplasm resources in cultivation from the point of rhizospheric bacteria.

## MATERIALS AND METHODS

### Experimental design

One of the main producing region of *A. dahurica* var. *formosana* is Suining in Sichuan province, China. In previous experiments, our research group has bred several varieties (strains) of *A. dahurica* var. *formosana* which were named as BZA001, BZA002, BZA003, BZA004, BZB002 and BZB003.

In order to explore the effects of different planting sites of the same varieties (strains) on rhizosphere bacteria of *A. dahurica* var. *formosana*, BZA001 was used as the test material, four different *A. dahurica* var. *formosana* planting sites in Suining and one in Chongzhou as the control with the same cultivation and management methods were investigated. The samples of five different experimental sites were named as XA (Shunjiang, Suining), XB (Shunhe, Suining), XC (Sangshulin, Suining), XD (Yongyi, Suining), XJ (Chongzhou, Chengdu). Meanwhile, in order to explore the rhizospheric bacteria of different varieties (strains) in the same site, six varieties (strains) of *A. dahurica* var. *formosana* were also sampled in one of sites in Suining (Yongyi, Suining), which were named as XD(BZA001), XE(BZA002), XF(BZA003), XG(BZA004), XH(BZB002) and XI(BZB003), respectively. All the *A. dahurica* var. *formosana* experimental materials were sown in late September, 2018. 750 kg/hm of superphosphate and 120 kg/hm of potassium phosphate were applied as base fertilizer. In December, the first topdressing application of urea was 66 kg/hm. The second topdressing application of urea 165 kg/hm was carried out in February, 2019. The third topdressing was carried out in March, 2019, applying urea 99 kg/hm, superphosphate 750 kg/hm and potassium sulfate 8 kg/hm. Plots within the field were established in randomized blocks design with each plot consisting of four rows (5 m long, 3 m wide) and three replicated plots per treatment.

All *A. dahurica* var. *formosana* samples were collected in harvest time (sown in September, 2018 and harvested in July, 2019). the five-point sampling method was used, representative and robust *A. dahurica* var. *formosana* were selected, the excess bulk soils were shaked off, and the soils that remained attached to the roots were considered to be rhizospheric soils (30). Soil samples were respectively preserved at −80 °C and 4 °C before used. Meanwhile, the roots of *A. dahurica* var. *formosana* were brought back after sampling the soil.

### Soil physicochemical properties

Soil samples were collected from the rhizosphere of *A. dahurica* var. *formosana* from different experimental sites or varieties (strains) for the determination of physicochemical properties. The soil pH was measured with a pHB-8 pen type acidity meterin in a 1:2.5 soil/water (W/V) suspension. The OM was measured by potassium dichromate volumetric method. Soil TN was determined by Kjeldahl method. Soil TP was determined by sodium hydroxide solution-Mo-Sb anti spectrophotometric method. Soil TK was measured by acid fusion-flame photometric method. The HN in the soil was determined by alkali hydrolysis diffusion method. The AP was obtained by using the sodium hydrogen carbonate solution-Mo-Sb anti spectrophotometric method. The available potassium was obtained by ammonium acetate extraction - flame photometric method.

### Yield and active component contents of *A. dahurica var. formosana*

Yield was estimated by the ratio of total fresh weight of root to area in each plot. Selected the representative root of *A. dahurica* var. *formosana* with the same growth trend, washed and water-removing at 105°C for 15 minutes, then dried at 55°C to a constant weight. The active components of *A. dahurica* var. *formosana* were coumarins. The contents of imperatiorin and isoimperatorin of *A. dahurica* var. *formosana* were determined respectively according to the Chinese Pharmacopoeia (2020 edition, part I) (6).

### Soil DNA extraction and MiSeq sequencing

Genomic DNA of soil samples were extracted by method of CTAB or SDS. The DNA concentration and purity were detected by agarose gel electrophoresis, and then an appropriate amount of samples were diluted with sterile water to 1ng/μL. The diluted genomic DNA was used as the template. The primers with Barcode for the 16S V3-V4 region primers were 341F: CCTAYGGGRBGCASCAG, and 806R: GGACTACNNGGGTATCTAAT. PCR amplification was carried out using Phusion ^®^ High-Fidelity PCR Master Mix with GC Buffer of New England Biolabs and high-fidelity enzyme to ensure the amplification efficiency and accuracy.

PCR products were detected by electrophoresis with 2% agarose gel. PCR products were mixed in equal amounts according to concentration. PCR products were purified by agarose gel electrophoresis with 1×TAE concentration of 2%, and the target bands were cut and recovered. The product purification kit USES GeneJET gel recovery kit to recover the product from Thermo Scientific company. Ion Plus Fragment Library Kit 48 rxns from Thermofisher was used for Library construction. After Qubit quantification and Library detection, Thermofisher Ion S5TMXL was used for computer sequencing. Raw reads were obtained from the reads by using Cutadapt (1) to shear low quality sequences and preliminary quality control. Raw Reads were detected by comparison with the species annotation database and chimeric sequences removal (26) and filtered to obtain Effective Tages.

### Statistical analysis

Sequences of clean reads with ≥97% similarity were assigned to the same OTUs (Operational Taxonomic Units). According to the algorithm, the sequence with the highest frequency was selected as the OTU representative sequence. According to the algorithm, the sequence with the highest frequency was selected as the OTU representative sequence. Species annotation analysis was carried out by Mothur method and SSUrRNA database (the threshold value was set as 0.8~1), acquire information taxonomy and respectively in each classification level: kingdom, phylum, class, order, family, genus, species counted the community composition of each sample (9, 10, 40). The samples were homogenized with the least amount of data as the standard. The Observed-OTUs, Chao1, Shannon, Simpson, ACE, Goods-coverage, PD whole tree were calculated by QIIME (Version 1.9.1). R (Version 2.15.3) was used to analyze the differences between groups of alpha diversity index and draw PCoA chart. In the analysis of species differences between groups, LEFSE software was used for LEFSE analysis. The filter value of LDA Score was set as 4 by default, and software R was used for T-test. In Spearman correlation analysis, Corr.test function in R was firstly used to calculate Spearman correlation values of species and environmental factors and test its significance, and then heatmap function was used for visualization.

## RESULTS

### The physicochemical properties of rhizospheric soil of *A. dahurica* var. *formosana*

Overall, shown in Table 1, the rhizospheric soil nutrients in Chongzhou were significantly higher than that in Suining from different experimental sites. The rhizospheric soil pH value of Chongzhou was neutral (7.1), and the others from Suining were alkaline (pH value from 7.8-8.0). Previous results showed that the pH of *A. dahurica var. formosana* soil in Suining producing area was about 8.49-8.88 (43), indicating that the rhizosphere soil of *A. dahurica var. formosana* was acidified, and the pH of soil was about 0.49-1.08 lower than that of non-rhizosphere soil. The organic matter (OM) content of Chongzhou rhizospheric soil is triple to tenfold that of other rhizospheric soils in Suining, up to 22.7 g/kg. OM of Suining’s rhizosphere soil varied from 5.04 to 7.34 g/kg. The rhizosphere soil total nitrogen (TN) value varied greatly, showing a 4.48-fold change (0.33-1.48 g/kg). Chongzhou had the highest TN (1.48 g/kg), while Yongyi of Suining had the lowest TN (0.23 g/kg). The rhizosphere soil total phosphorus (TP) value varied from 0.52 to 0.98g/kg, with the highest in Shunjiang (0.98 g/kg) and Shunhe (0.96 g/kg), and the lowest (0.52 g/kg) in Chongzhou. Rhizosphere soil total potassium (TK) had a small variation range (13.41-16.15 g/kg), with the highest (16.15 g/kg) in Shunhe and the lowest (13.41 g/kg) in Yongyi. The rhizosphere soil hydrolysable nitrogen (HN) value changed greatly, showing a 4.60-fold change (17.5-80.50 mg/kg). The HN of rhizospheric soil of *A. dahurica var. formosana* from Chongzhou was the highest (80.50 mg/kg), while that of Yongyi was the lowest (17.50 mg/kg). The rhizosphere soil available phosphorus (AP) value changed greatly, showing a 4.00-fold change (4.42-17.69 mg/kg), with the highest from Chongzhou (17.69 mg/kg), and the lowest from Yongyi (4.42 mg/kg). The rhizospheric soil available potassium (AK) value changed greatly from 14.18 to 195.62mg/kg with the highest from Chongzhou (195.62 mg/kg) and the lowest from Sangshulin (14.18 mg/kg).

**TABLE 1.**
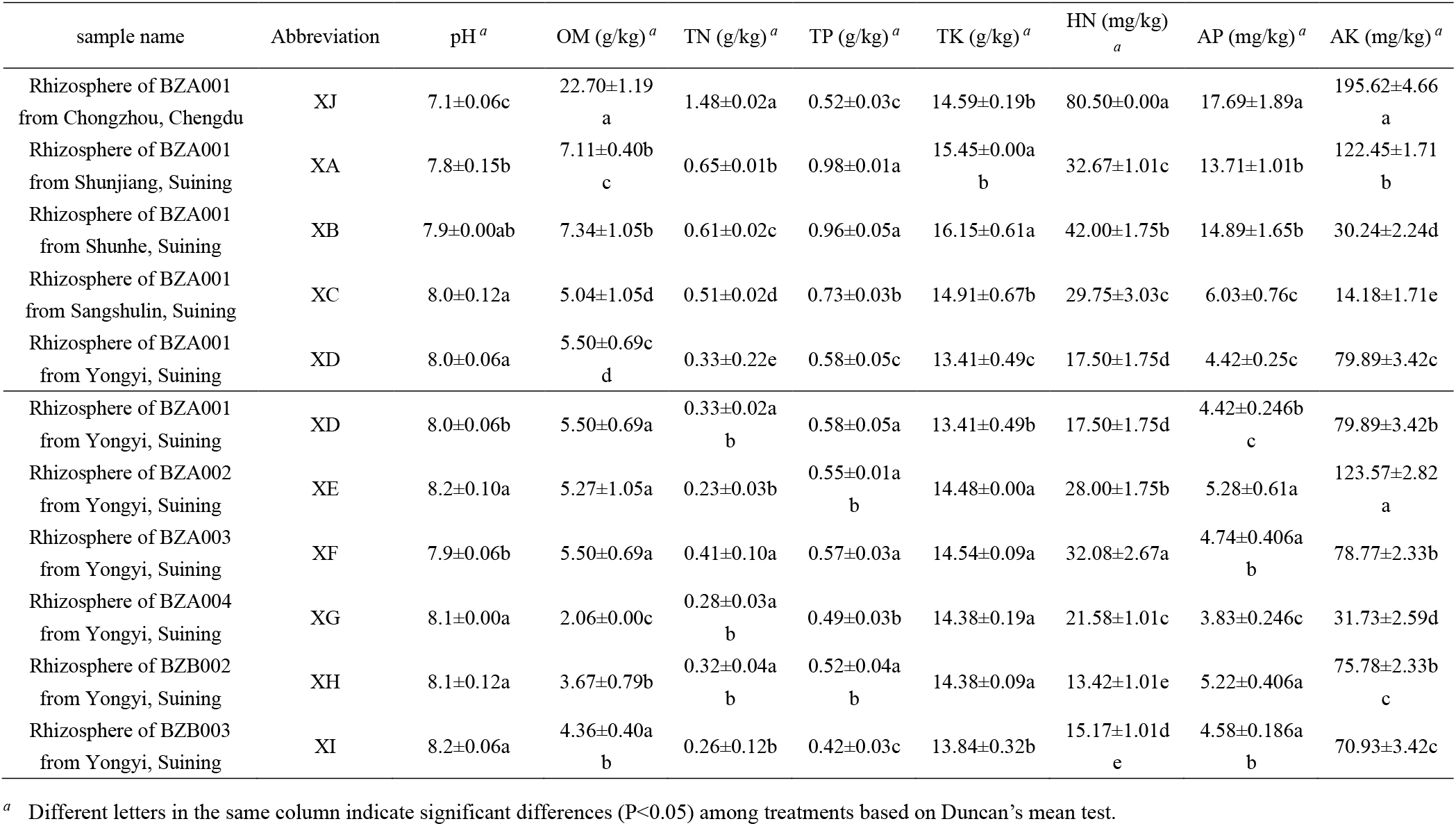
The physicochemical properties in the rhizospheric soil of *A. dahurica var. formosana* from different experimental sites and varieties (strains).

Among all varieties (strains), overall, BZA002 had the highest rhizosphere nutrients of *A. dahurica* var. *formosana*, while BZA004 had the lowest. The pH value changed little (7.9-8.2). The OM of Yongyi varied from 2.06-5.50 g/kg, among which BZA004 had the lowest (2.06 g/kg). The TN value changed little (0.23-0.41 g/kg). The TP value varied from 0.42 to 0.58g/kg, of which BZB003 was the lowest (0.42 g/kg). TK had a small variation range (13.41-14.54 g/kg). The HN value of rhizospheric soil was low (13.42-32.08 mg/kg), of which BZA003 was the highest (32.08 mg/kg) and BZB002 was the lowest (13.42 mg/kg). The AP value was low (3.83-5.28 mg/kg), too. The rhizospheric soil AP of BZA002 was the highest (5.28 mg/kg), while that of BZA004 was the lowest (3.83 mg/kg). The AK value changed greatly from 31.73 to 123.5 mg/kg, of which BZA002 was the highest (123.57 mg/kg) and BZA004 was the lowest (31.73 mg/kg).

### The yields and active component contents of *A. dahurica var. formosana* from different experimental sites and varieties (strains)

For different experimental sites of the same varieties (strains) of *A. dahurica var. formosana*, the yield in Suining was about 3-4 times that of Chongzhou, although the active component contents of *A. dahurica var. formosana* of Suining were not as high as that of Chongzhou (Table 2). To be specific, the yield of *A. dahurica var. formosana* from Yongyi was the highest (33603.46kg/hm), and that from Chongzhou was the lowest (9171.25kg/hm). The imperatorin content of *A. dahurica var. formosana* in Chongzhou was the highest (2.83 mg/g), and that of Yongyi was the lowest (1.50 mg/g). The isoimperatorin content in Shunjiang was the highest (1.34 mg/g), followed by Chongzhou (1.03 mg/g), and Yongyi was the lowest (0.85 mg/g).

**TABLE 2.**
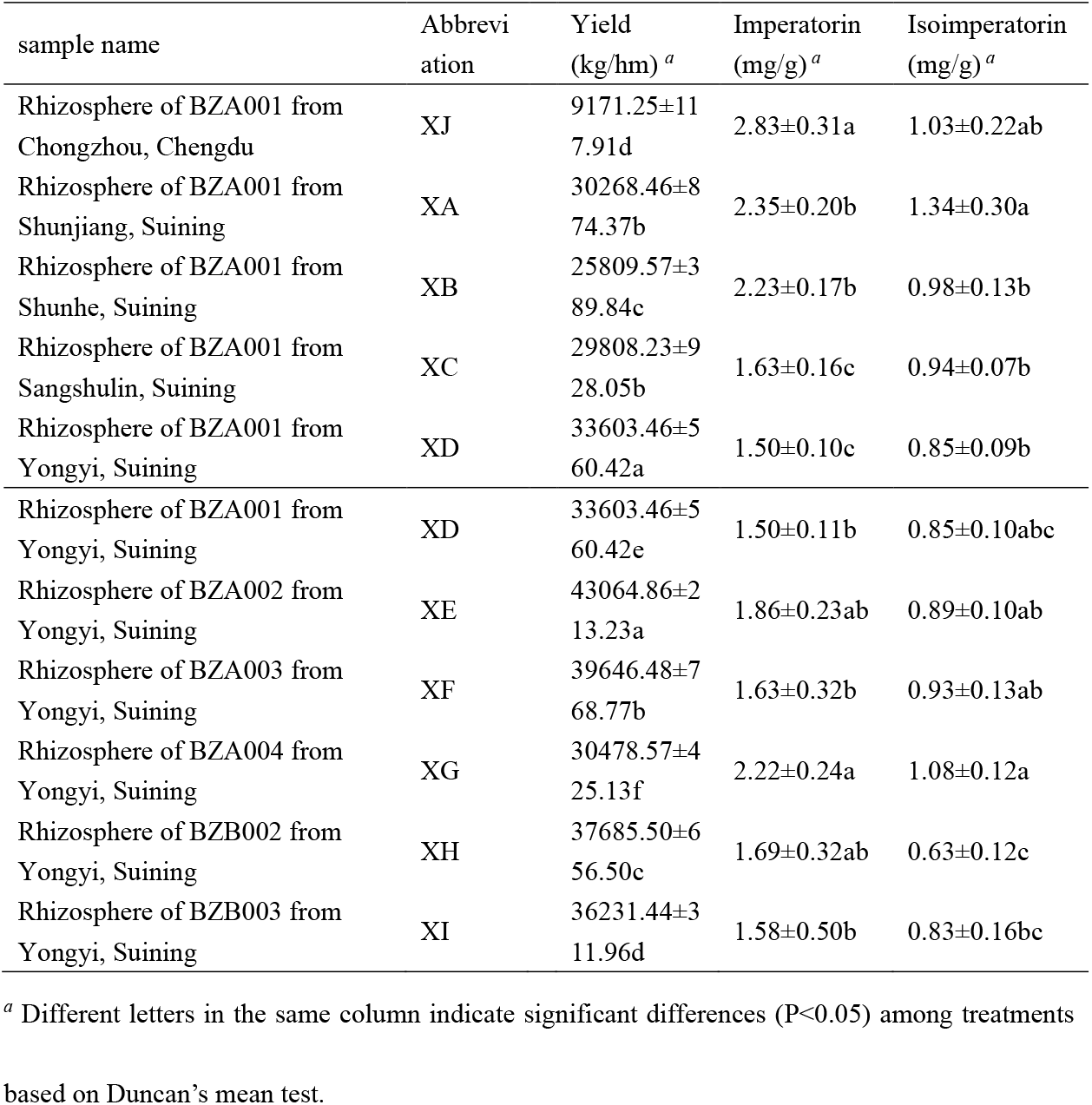
The yields and active component contents of *A. dahurica var. formosana* from different experimental sites and varieties (strains)

For different varieties (strains) of *A. dahurica var. formosana* from the same experimental sites, the yield of BZA002 was the highest (43064.86 kg/hm) and that of BZA004 was the lowest (30478.57 kg/hm). The imperatorin content of BZA004 was the highest (2.22 mg/g), and that of BZA001 was the lowest (1.50 mg/g). The isoimperatorin content of BZA004 was the highest (1.08 mg/g) and BZB002 the lowest (0.63 mg/g).

### Bacterial diversity in the rhizospheric soil of *A. dahurica var. formosana*

Through comparison with database SILVA132, species annotation and statistics of different classification levels, a total of 16,557 OTUs were found, of which 16,557 (100.00%) could be annotated to the database. As shown in Table 3, for the same varieties (strains), the rhizosphere bacterial diversity of *A. dahurica var. formosana* in Chongzhou experimental site with high nutrients and low yield was the lowest, while the other four experimental sites in Suining were significantly higher. Among them, the rhizosphere bacterial diversity from Yongyi with the lowest nutrients was the highest. For different varieties (strains) of *A. dahurica var. formosana*, the rhizospheric bacterial diversity of BZA002 with high nutrients and yield was less than that of the others from Yongyi, while the bacterial diversity of BZB003 was the highest. Spearman correlation analysis with soil nutrients showed that, except for AK, the rhizosphere diversity of *A. dahurica var. formosana* was significantly negatively correlated with soil nutrients (Fig S1).

**TABLE 3.**
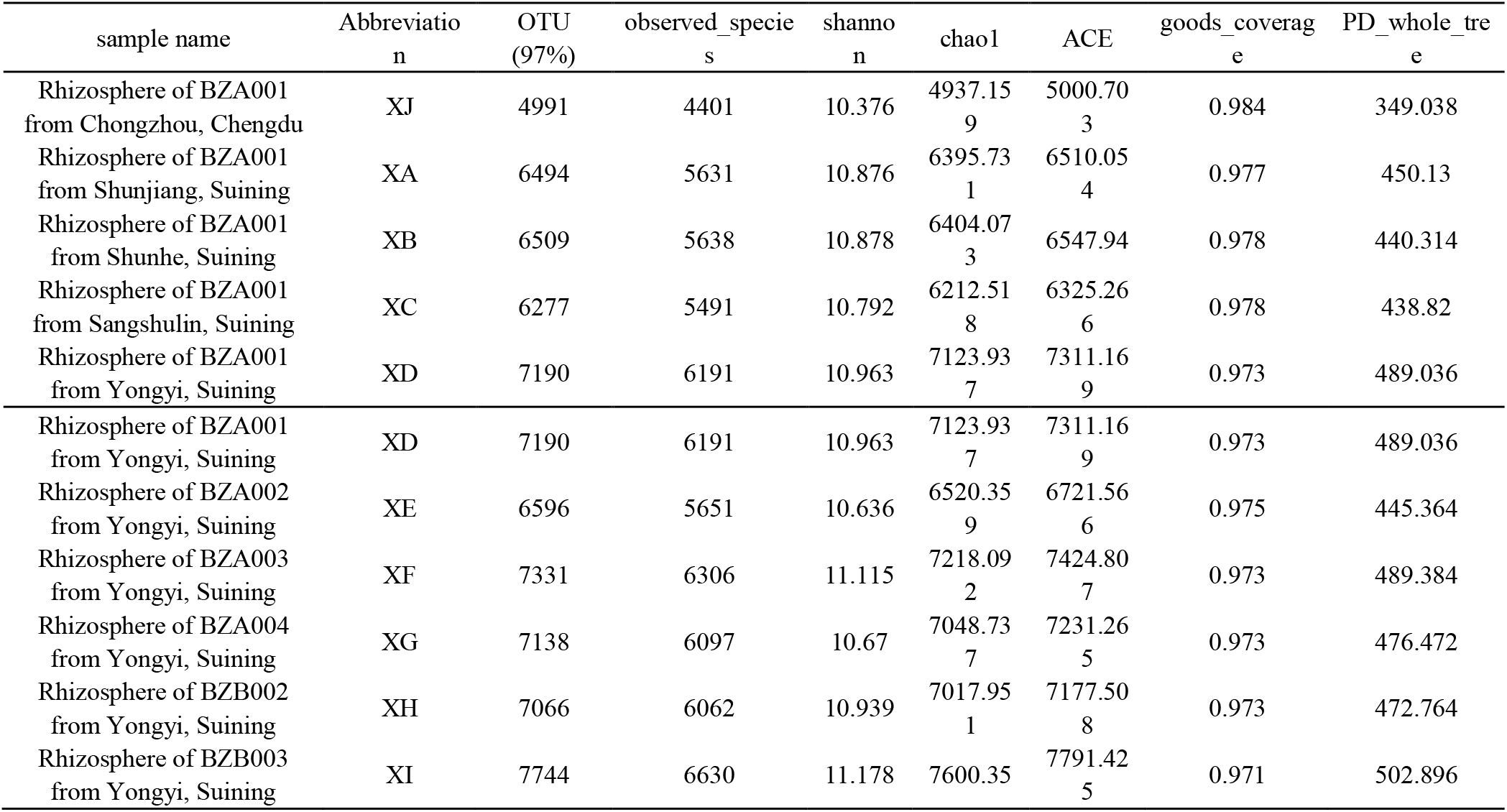
Diversity indices of the bacterial community in rhizospheric soil samples of *A. dahurica* var. *formosana* from different experimental sites and varieties (strains).

### Relative abundances of rhizospheric bacterial communities in *A. dahurica var. formosana*

In the rhizosphere soil of *A. dahurica var. formosana, Proteobacteria, Firmicutes* and *Bacteroidetes* were the dominant phylum (Fig 1). For different experimental sites, sample from Yongyi (XD) had the highest levels of *Proteobacteria* sequences(49.1%) and the lowest *Acidobacteria*(6.6%) and *Rokubacteria* sequences(1.1%). The proportions of *Firmicutes* and *Actinobacteria* in the rhizosphere soil samples of *A. dahurica var. formosana* from Chongzhou were significantly higher than those in other Suining sites, showing ratios of 7.2% and 8.1%. For different varieties(strains) of *A. dahurica var. formosana* from Suining, the proportion of *Proteobacteria* was highest in the rhizosphere of BZA002 and lowest in the rhizosphere of BZA004. The proportion of *Acidobacteria* was highest in the rhizosphere of BZA002 and lowest in the rhizosphere of BZB003. The proportion of *Actinobacteria* was lowest in the rhizosphere of BZA001. The proportion of *Verrucomicrobia* was lowest in the rhizosphere of BZA004.

**FIG. 1.**
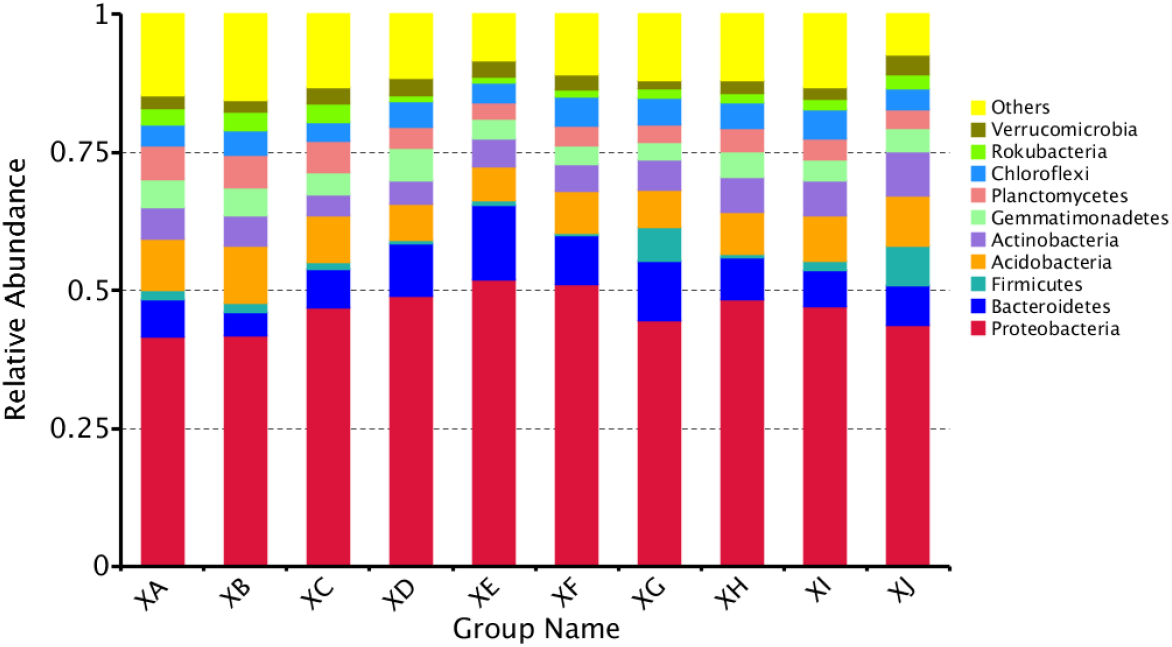
Relative species abundance at phylum level in bacterial communities (A) from different sites and (B) from different varieties (strains). Based on the species annotation results, we selected to examine the relative abundance distribution of the species among the top-ranked phyla, the relative abundance distribution of the top ten species is shown.

At the genus level, we selected the 35 of the most abundant genera to generate a mapping image (Fig. 2). The abundance of the sorting results, ordered from high to low, were *Bacteroides*, *Flavobacterium*, unidentified *Clostridiales*, *Anaeromyxobacter*, *Sphingomonas*, *Faecalibacterium*, *Polycyclovorans*, *unidentified Acidobacteria*, *Haliangium*, *Terrimonas*, etc. *Anaeromyxobacter*, *Sphingomonas*, *Polycyclovorans* and *Haliangium* belonged to the phylum *Proteobacteria*. A higher proportion of the genera *Haliangium*, *Anaeromyxobacter* and unidentified *Clostridiales* was detected in the rhizospheric bacteria of BZA001 from Chongzhou (XJ) with the highest nutrients. *Sphingomonas*, *Flavobacterium* and *Terrimonas* accounted for a higher proportion among the high yield BZA002 rhizosphere bacteria (XE). *Faecalibacterium* and *Bacteroides* accounted for a higher proportion among the high yield BZA004 rhizosphere bacteria (XG).

**FIG. 2.**
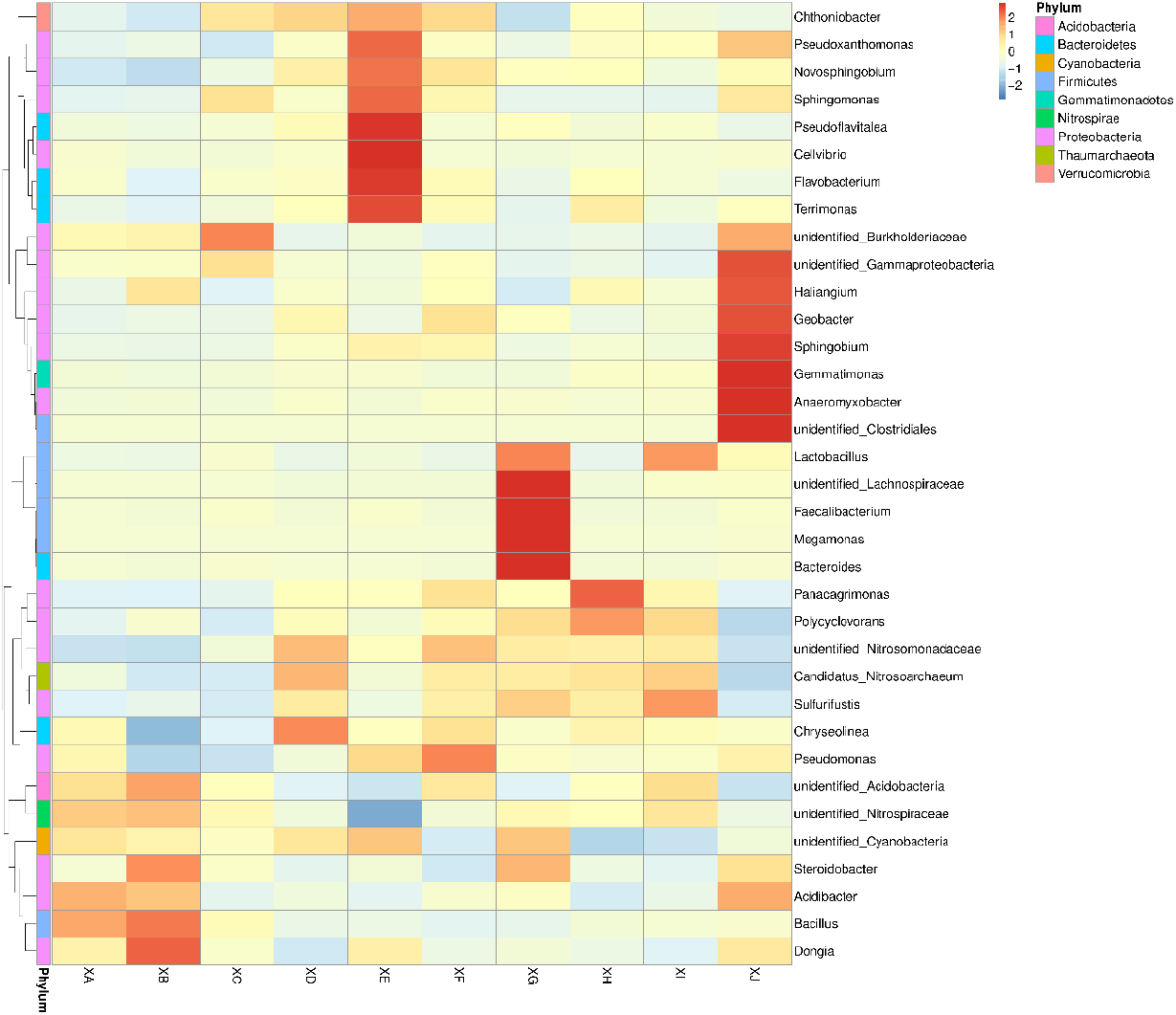
Generate a mapping image from different sites and from different strains based on abundance in formation. To examine the abundance information available, we selected genera represented by the 35 most abundant sequences to draw a species abundance cluster heatmap. Abscissae indicate species annotation information, and ordinates indicate sample information. Clustering tree on the left side of the figure is a sample cluster tree. Above the figure is a species tree clustering and the results of the significance test of the species in different groups.

### PCoA cluster analysis in the rhizospheric soil bacteria of *A. dahurica var. formosana*

Comparison of PCoA among treatment groups was conducted to explore the similarity and difference of community composition among different groups. The results showed that there were significant differences among all groups (Fig 3, Table S1). For different experimental sites of the same varieties (strains) of *A. dahurica var. formosana*, the explanatory variances of principal component 1 and principal component 2 were 27.85% and 15.79% respectively. The rhizosphere bacterial community of *A. dahurica* var. *formosana* in Chongzhou (XJ) had the largest difference, followed by that in Yongyi of Suining (XD). The rhizosphere bacterial community of Shunjiang (XA), Shunhe (XB) and Sangshulin (XC) experimental sites were similar. For different varieties (strains) of *A. dahurica var. formosana* from the same experimental sites, the explanatory variances of principal component 1 and principal component 2 were 10.57% and 8.56%, respectively. The rhizospheric bacterial communities of BZA002 and BZA004 were the most different from those of other varieties (strains). The results indicated that there were significant differences among the experimental groups in the selected experimental sites and the background of varieties (strains). Moreover, there were significant differences between Suining and Chongzhou. Among them, the dominant factors of experimental sites affecting the rhizosphere bacterial community structure of *A. dahurica var. formosana* were more obvious.

**FIG 3.**
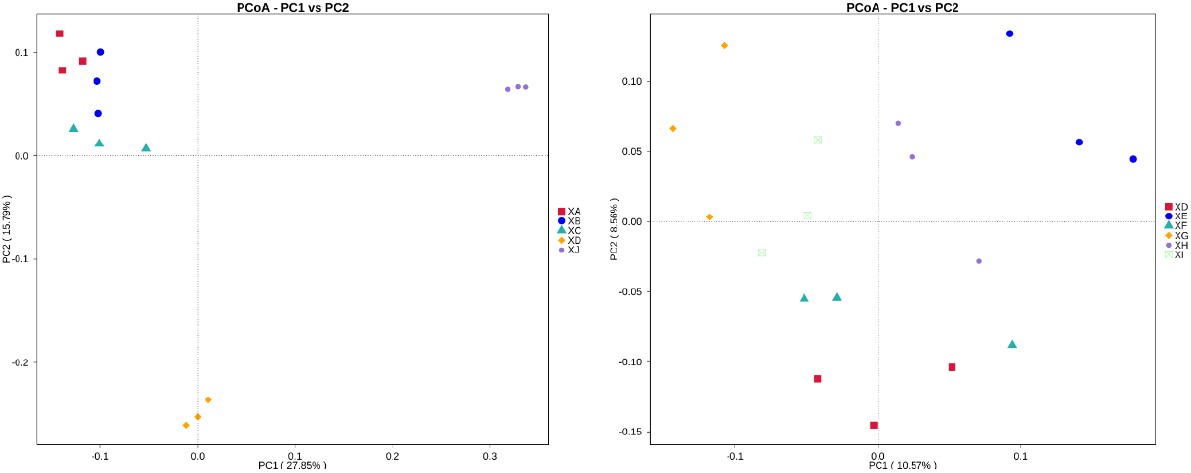
Principal co-ordinates analysis (PCoA). The main elements and structures were extracted from the multi-dimensional data by eigenvalue and eigenvector sorting, and the main coordinate combination with the largest contribution rate was selected for drawing and display.

### LEfSe analysis in the rhizospheric soil bacteria of *A. dahurica var. formosana*

The biomarkers with significant differences in the rhizosphere samples of *A. dahurica var. formosana* from different sites or different varieties (strains) through LEfSe analysis were revealed (Fig 4), and linear discrimination analysis (LDA) values were given in Fig 5.The results showed that at the phylum level, for different experimental sites, enrichment of *Planctomycetes* were significant in the rhizosphericsoil of BZA001 from Shunjiang (XA); enrichment of *Acidobacterira* were significant in the sample from Shunhe (XB); enrichment of *Roteobacteria* were significant in the sample from Sangshulin (XC); *Proteobacteria* contributed the most in the sample from Yongyi (XD) with the highest yield of *A. dahurica var. formosana*, while *Firmicutes* and *Actinobacteria* contributed the most in the sample from Chongzhou (XJ) with the highest active component content of *A. dahurica var. formosana*. There were 21 biomarkers with an LDA score >4, including *Gammaproteobacteria*, *Anaeromyxobacter*, *Sphingomonadales*, *Rokubacteria*, *Longimicrobiales*, *Firmicutes*, etc (Fig 4A, 5A). These biomarkers could be considered to represent the key bacterial taxa shaping the rhizosphere bacterial community of *A. dahurica var. formosana*.

**FIG 4.**
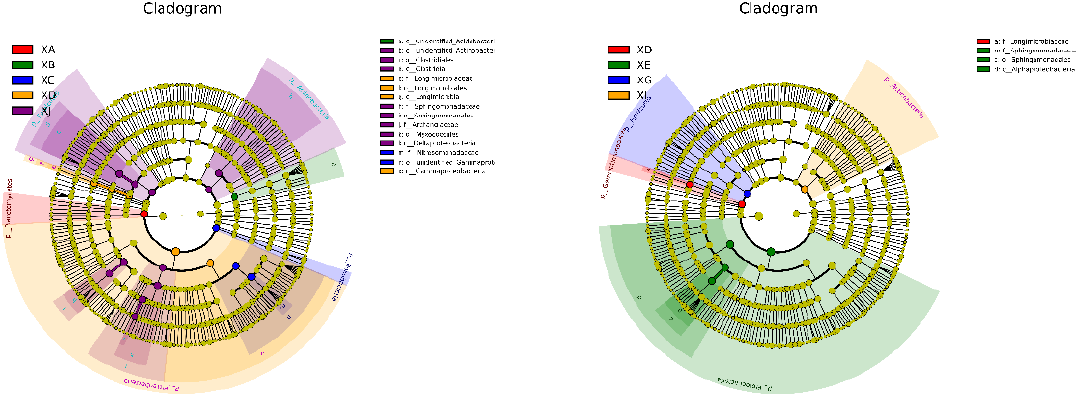
Linear discriminant analysis effect size (LEfSe) method identifies the significantly different abundant taxa (a) from different experimental sites and (b) from different varieties (strains) in all of the rhizospheric soil samples of *A. dahurica var. formosana*. The groups with significant differences in relative abundance among treatments were represented by colored dots, which represented the level of boundary, phylum, class, order, family and genus from the center outward. The size of the colored dots was proportional to the relative abundance. The biomarker of different species followed the group for staining, represented by “a:”, “b:”, “c:”, etc.

**FIG 5.**
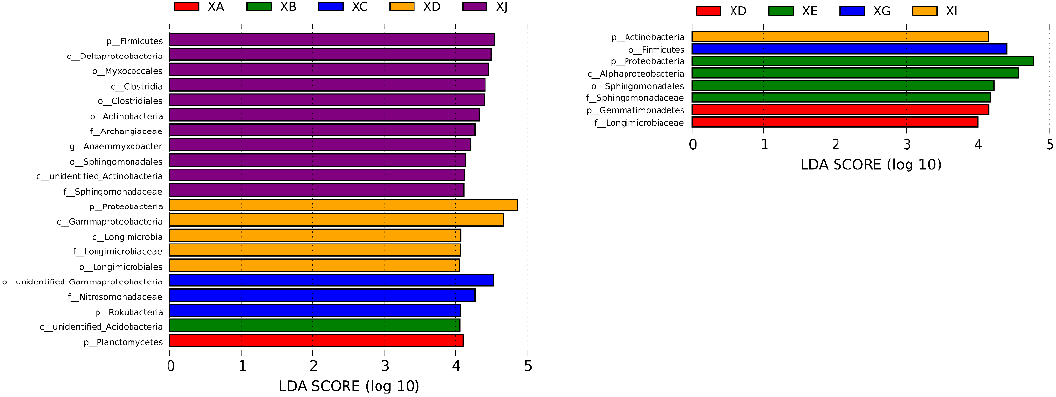
The LDA scores of the soil from different experimental sites(a) and different varieties (strains) (b) in each treatment from the LEfSe analysis. The figure shows species with significant difference in different (LDA value greater than 4.0 species with difference), and the length of the bar chart represents the impact of significantly different species.

From the same experimental sites, the major divergent species of different varieties (strains) of *A. dahurica var. formosana* rhizospheric soil were also differed. For example, *Gemmatimonadetes* contributed the most in the bacterial community of BZA001 rhizospheric soil (XD); *Proteobacteria* contributed the most in the bacterial community of BZA002 rhizospheric soil (XE) with the highest yield of *A. dahurica var. formosana*; *Firmicutes* contributed the most in the bacterial community of BZA004 rhizospheric soil (XG) with the highest active component content of *A. dahurica var. formosana*; *Actinobacteria* contributed the most in the bacterial community of BZB003 rhizospheric soil (XI). There were 8 biomarkers with an LDA score>4, including *Sphingomonadales*, *Firmicutes*, *Alphaproteobacteria*, *Longimicrobiaceae*, *Actinobacteria*, *Proteobacteria*, etc (Fig 4b, 5b).

Combined with the results of BZA001 rhizospheric bacterial from five experimental sites, it was found that *Proteobacteria* had a significant influence on the rhizosphere bacterial community of the samples with higher yield, and *Firmicutes* had a significant influence on the rhizosphere bacterial community of the samples with higher content of active components, regardless of whether the influencing factors were from the differences of experimental sites or varieties (strains). In addition to the phylum level, Sphingomonadaceae was found to be significantly enriched in soils with high nutrients and low α diversity.

### Correlation analysis between environmental physicochemical factors and bacterial taxa

The results of spearman correlation heatmap analysis showed that, in general, the physicochemical properties of rhizospheric soil (except for AK) were closely related to bacterial taxa, especially AP and TN exerted the greatest impacts upon microbial community composition (Fig 6). The abundance of *Acidobacteria* (r=0.5694) and *Rokubacteria* (r=0.6158) were most closely related to AP, and that of *Acidobacteria* (r=0.6209) and *Rokubacteria* (r=0.6653) were also most closely related to TN. Moreover, the abundance of *Proteobacteria* was significantly negatively correlated with AP(r=-0.4811) and TN (r=-0.5790), and abundance of *Firmicutes* was significantly positively correlated with HN (r=0.4895) and TN (r=0.4927). The abundance of *Actinobacteria* was only negatively correlated with TP (r=-0.3783), while the abundance of *Verrucomicrobia* was positively correlated with AK (r=0.4758).

**FIG 6.**
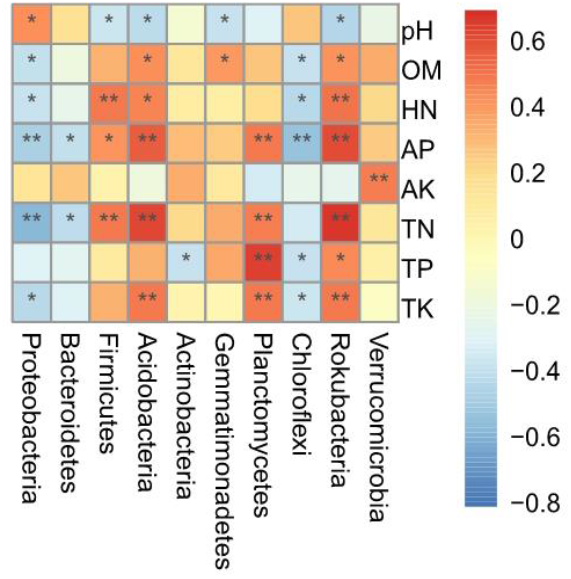
Spearman correlation heatmap analysis of the correlation between soil factors and species abundance of bacterial communities in rhizospheric soil of *A. dahurica var. formosana* at phylum level. The vertical is environmental factor information, and the horizontal is species abundance information. The value corresponding to the middle heat map is Spearman correlation coefficient r, which is between −1 and 1. “*” indicates significant difference (P<0.05).

## Discussion

Analysis of microbial diversity and community composition using high-throughput sequencing technology is becoming more and more common (24). To our knowledge, more and more studies had been conducted on the interaction between medicinal plants and their rhizosphere microorganisms, especially those that could improve the yield and quality of medicinal plants (16, 17, 18, 34, 41). Thus, this new sequencing technique was used for the first time to analyze the bacterial community structure in the rhizosphere soil of *A. dahurica var. formosana*.

It was possible that *A. dahurica* var. *formosana* recruited less beneficial bacteria when the soil was rich in nutrients and more bacteria when it was hungry based on the correlation analysis of bacterial diversity and physicochemical properties of rhizosphere samples. In this study, the content of active components of *A. dahurica var. formosana* in each group was far higher than the industry standard. The yield of *A. dahurica var. formosana* in Suining is 3-4 times that of Chongzhou, although the content of active components in *A. dahurica var. formosana* in Suining is lower than that in Chongzhou. By comparison, *A. dahurica var. formosana* from Suining, the traditional main producing area, is still the best, while its rhizosphere nutrient content is lower. Mutualism exists between plants and a variety of microorganisms and provides more than half of a plant’s nutritional needs (14). In this experiment, except AK, soil physicochemical properties were negatively correlated with bacterial diversity. Chaudhary found that rhizospheric bacteria could significantly improve the utilization rate of soil N, P and K, replacing the deficiency of soil nutrients to a certain extent (5). To our knowledge, when the phosphorus content in soil was too low, legumes would stimulate the enrichment of AMF in the rhizosphere by secreting isoflavones to promote nutrient absorption (23). Meanwhile, when soil phosphorus content is too high, plants can obtain the optimal nutrient supply without AMF symbiosis instead of preferring symbiosis (11). Therefore, ensuring bacterial diversity in the rhizosphere of *A. dahurica var. formosana* may be a new idea for fertilizer reduction.

There may be some different secretions in the root of BZA002, which activated the nutrient availability of rhizospheric soil and promotes the growth of *A. dahurica* var. *formosana* more than other varieties (strains). At the same experimental site, the soil nutrient availability was the highest in the rhizosphere of BZA002 with highest yield, and the lowest in the rhizosphere of BZA004 with lowest yield. Such differences in rhizosphere soil nutrients are caused by differences in germplasm. Different varieties secrete different root exudates, which lead to differences in recruitment of beneficial bacteria and nutrient uptake by the root system (44). Yang found that there were significant differences in the activation ability of soil K by tobacco root exudates with different genotypes (42). Li pointed out that compared with the general genotype, the root exudates of grain amaranth with potassium-enrichment genotype had higher organic acid content, which could significantly activate the mineral potassium in the soil (21). Shen found that kidney beans with different genotypes could secrete plenty of organic acids to activate Al-P in the rhizosphere under P deficiency conditions (15).

Imperatorin might be one of the reasons for the decrease of rhizosphere microbial diversity of the highly active ingredient *A. dahurica var. formosana*. The main active components in *A. dahurica var. formosana* were coumarins and essential oils, and coumarins mainly include imperatorin and isoimperatorin. It was found that regardless of the difference of origin or germplasm, the variation trend of imperatorin was inverse direction to that of bacterial diversity. To our knowledge, Coumarins in *A. dahurica var. formosana* had antibacterial activity, which could recruit more phosphate-solubilizing microorganisms (32) and had inhibitory effect on a variety of pathogens (27).

*Proteobacteria*, *Firmicutes* and *Bacteroidetes* were dominant phylum of *A. dahurica var. formosana*, accounting for 47.6%-66.4% of all bacterial DNA sequences. These eutrophic bacteria can mineralize soil nutrients, enhance the absorption of plant nutrients, and promote plant growth (36). This is similar to the dominant rhizospheres of healthy *Panax notoginseng* and licorice (35, 45). Regardless of the difference of experimental tests or varieties (strains), *Proteobacteria* were all the dominant phyla in the rhizosphere of *A. dahurica var. formosana*, and the dominant species at the genus level were also similar. The results indicated that the rhizosphere bacterial community of *A. dahurica var. formosana* was stable and conserved, and could resist some external disturbances to a certain extent, which provided a theoretical basis for remote planting and expanding the planting.

*Acidobacteria* and *Rokubacteria* were prevalent in the rhizosphere of *A. dahurica var. formosana*. There were a variety of gene clusters for the synthesis of antibiotics, antifungins, ferritin and immunosuppressants in these bacteria (7). In the similar environment of the four experimental sites in Suining, the bacterial diversity (XD) was high when the content of *Acidobacteria* and *Rokubacteria* in the rhizosphere was low, suggesting that these bacteria with antibacterial activity might inhibit the colonization of other bacteria in the rhizosphere of *A. dahurica var. formosana*. Due to continuous cropping obstacles, the rhizosphere microflora of *P. notoginseng* and other peanut species changed from “bacterial-type” to “fungal-type” during continuous cropping period, and the bacterial community of Proteobacteria and other bacteria showed a downward trend (22, 35). No significant continuous cropping obstacles were found in *A. dahurica var. formosana*, which might be due to its antibacterial active components and bacterial community, which inhibited the occurrence of soil-borne diseases.

*Proteobacteria* and *Firmicutes* might play a role in increasing yield and active ingredient content. By LEfSe analysis of five experimental sites (XA-XB-XC-XD-XJ), we found that there were 21 taxa of differential distributions with LDA score >4. Among these biomarkers, *Proteobacteria* enrichment in the *A. dahurica var. formosana* rhizospheric soil of high-yield site (XD) was significant, while *Firmicutes* and *Actinobacteria* enrichment in the rhizospheric soil of high-active components of *A. dahurica var. formosana* (XJ). Therefore, *Proteobacteria*, *Firmicutes* and *Actinobacteria* might be closely related to yield and content in terms of origin factors. By LEfSe analysis of six varieties (strains) of *A. dahurica var. formosana* (XD-XE-XF-XG-XH-XI), there were 8 taxa of differential distributions with LDA score >4. *Proteobacteria* is significantly enriched in the rhizosphere of high-yield BZA002. *Firmicutes* was significantly enriched in the rhizosphere with high content of active components of BZA004. *Actinobacteria* was significantly enriched in the rhizosphere with low-active components of BZB003. In general, *Proteobacteria* were always significantly enriched in the rhizosphere of the high-yield samples, while *Firmicutes* were always significantly enriched in the rhizosphere of the high-active component of *A. dahurica var. formosana*, regardless of the difference of experimental tests or varieties (strains).

Interestingly, *Sphingomonadaceae* were found in differential taxas with LDA score>4 for soil environmental differences or genotypic differences. *Sphingomonadaceae* was significantly enriched in the samples with the highest soil nutrients and the lowest bacterial diversity. Many strains of *Sphingomonadaceae* can degrade polycyclic or monocyclic aromatic compounds, such as benzoic acid salicylic acid, and use these aromatic compounds as carbon sources for growth (31). Lin believed that allelopathic rice could secrete more aromatic compounds to induce the enrichment of these bacteria under chemical chemotaxis (20). *A. dahurica var. formosana* contained a large number of aromatic compounds. Similarly, a similar allelopathy may exist in *A. dahurica var. formosana*. The increasing of active components in *A. dahurica var. formosana* caused more aromatic compounds to be secreted from the root, recruiting more *Sphingomonadaceae* and anti-bacteria. Further studies on root exudates of *A. dahurica var. formosana* should be carried out to verify this hypothesis.

Understanding the key factors affecting the rhizosphere bacterial construction of *A. dahurica var. formosana* is an effective method to actively regulate the establishment of beneficial rhizosphere bacterial community, improve the rhizosphere environment and promote plant growth. As we know, in addition to suitable cultivation and management methods, good germplasm and suitable environment were crucial influencing factors for good medicinal plants. The rhizosphere bacterial communities of plants were also affected by these factors. Assembly of the rhizosphere microbial community was controlled by a complex interaction between microorganisms, plant hosts, and the environment (37). Among these factors, the stronger the environmental screening effect of which factor is, the greater the relative contribution to microbial community construction is likely to be (4).

The suitable origin and germplasm did have significant influence on the construction of rhizosphere bacterial community of *A. dahurica var. formosana*. PCoA indicated that the rhizosphere bacterial community of *A. dahurica var. formosana* showed significant differences among different experimental sites and varieties (strains). This confirmed the importance of suitable origin and germplasm for cultivation of medicinal plants from the perspective of rhizosphere bacteria, which was of great significance for improving the yield and quality of *A. dahurica var. formosana*.

Compared with germplasm resources, suitable planting site was the key factor for the construction of rhizosphere microbial community of *A. dahurica var. formosan*, which was of great significance for future development and application to improve the yield and quality. The four sites of Suining were all located along the coast of the Fujiang River and shared the similar climate. Excepted for the Chongzhou test site (XJ) with different climate, the differences among experimental sites were still slightly greater than the differences among varieties (strains). The results indicated that the rhizosphere bacterial community of *A. dahurica var. formosana* was more affected by soil in this study. Previous studies had shown that for many plants, genotype had a significant influence on the bacterial community in the aboveground part, especially perennial plants, while the rhizospheric bacterial community is mostly affected by soil factors (38). The difference of sorghum varieties had little effect on the rhizosphere bacteria, while the soil was the main factor (29). Automated ribosomal intergenic spacer analysis (ARISA) was used to investigate 9 rice varieties treated with different nitrogen levels. It was found that the varieties had a significant effect on the bacterial community of the stem and bud of rice, while the bacterial community of the root was mainly affected by the nitrogen level in the paddy field (28). Similarly, the rhizosphere bacterial community of *A. dahurica var. formosana* was significantly affected by soil of origin.

In addition to differences in soil nutrients, the original soil bacterial community might be another soil factor that affected the rhizosphere bacteria of *A. dahurica var. formosana*. When De Ridder-Duine studied the composition of rhizosphere and non-rhizosphere bacteria in different soils, he found that the structure of rhizosphere bacterial community was largely determined by the non-rhizosphere soil bacterial community (25). In other words, the original soil bacterial community played an important role in the construction of plant rhizosphere bacteria.

In general, this research results would be helpful in improving the cultivation and management of *A. dahurica var. formosana*. The results showed that the bacterial community of *A. dahurica* var. *formosana* was stable and conserved to a certain extent, and the bacteria of *Proteobacteria* and *Firmicutes* might play an important role in improving the yield and quality of *A. dahurica* var. *formosana*. The development and utilization of the rhizosphere microbial resources of *A. dahurica* var. *formosana* provided a new idea for further increasing production and quality and reducing the application of pesticide and fertilizer. The importance of germplasm resources and origin for the cultivation of *A. dahurica* var. *formosana* was discussed from the perspective of rhizosphere bacteria for the first time, among which the origin was the key factor affecting the composition of rhizosphere bacterial community. Soil conditions of origins would become a crucial factor to construct beneficial bacterial community structure in the cultivation of *A. dahurica* var. *formosana*, which laid a theoretical foundation for expanding the planting area in the future. However, further experiments were needed to explore the key influencing factors of soil factors, such as soil fertility, soil original microbiome, etc.

## Acknowledgment

This work was supported by Sichuan Science and Technology Program (No. 2021YFYZ0012) and Joint Implementation of Key R & D Projects in Sichuan and Chongqing in 2020(No. 2020YFQ0054).

## Appendixes

**TABLE S1.**
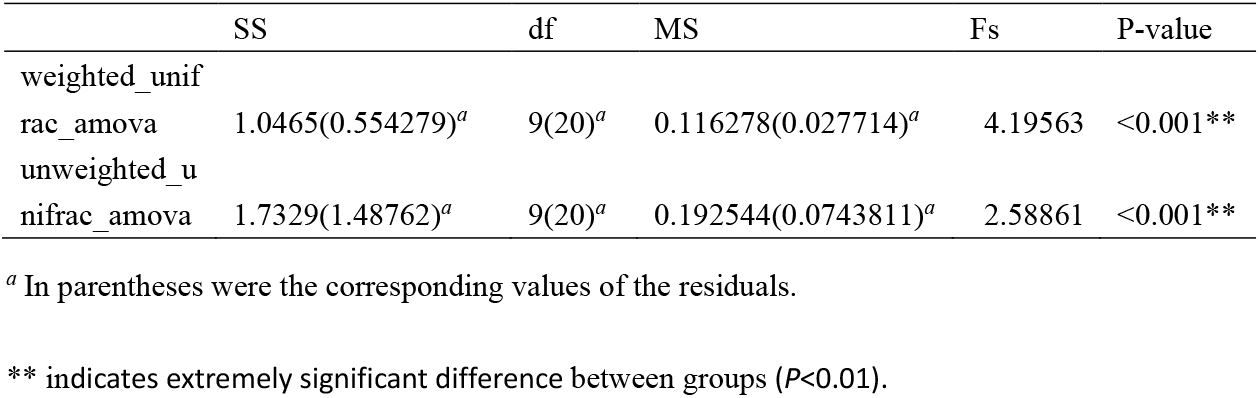
Analysis of molecular variance of rhizosphere bacterial community of *A. dahurica var. formosana*

**FIG S1.**
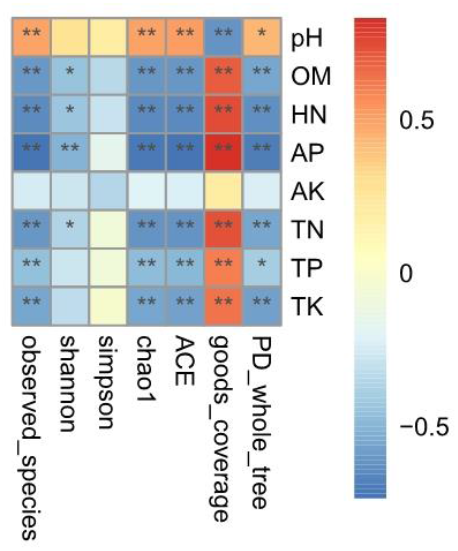
Spearman correlation heatmap analysis of the correlation between soil factors and diversity indices of bacterial communities in rhizospheric soil of *A. dahurica var. formosana*. The vertical is environmental factor information, and the horizontal is diversity index. The value corresponding to the middle heat map is Spearman correlation coefficient r, which is between −1 and 1. “*” indicates significant difference (P<0.05).

